# A Mutation-driven oncofetal regression fuels phenotypic plasticity in colorectal cancer

**DOI:** 10.1101/2023.12.10.570854

**Authors:** Slim Mzoughi, Megan Schwarz, Xuedi Wang, Deniz Demircioglu, Gulay Ulukaya, Kevin Mohammed, Federico Di Tullio, Carlos Company, Yuliia Dramaretska, Marc Leushacke, Bruno Giotti, Tamsin Lannagan, Daniel Lozano-Ojalvo, Dan Hasson, Alexander M. Tsankov, Owen J Sansom, Jean-Christophe Marine, Nick Barker, Gaetano Gargiulo, Ernesto Guccione

**Affiliations:** Center for OncoGenomics and Innovative Therapeutics (COGIT); Center for Therapeutics Discovery, Department of Oncological Sciences and Pharmacological Sciences, Tisch Cancer Institute, Icahn School of Medicine at Mount Sinai, New York, New York, USA; Department of Oncological Sciences, Icahn School of Medicine at Mount Sinai, New York, New York, USA; Graduate School of Biomedical Sciences at Icahn School of Medicine at Mount Sinai, New York, New York, USA; Tisch Cancer Institute Bioinformatics for Next Generation Sequencing (BiNGS) Shared Resource Facility, Icahn School of Medicine at Mount Sinai, New York, NY, USA; Max-Delbrück-Center for Molecular Medicine (MDC), Robert-Rössle-Str. 10, 13125 Berlin, Germany; Institute of Molecular and Cell Biology (IMCB), Agency for Science, Technology and Research (A*STAR), 61 Biopolis Drive, Proteos, Singapore 138673, Republic of Singapore; Department of Genetic and Genomic Sciences, Icahn School of Medicine at Mount Sinai, New York, New York, USA; Cancer Research UK Beatson Institute, Glasgow, UK; Department of Dermatology, Icahn School of Medicine at Mount Sinai, New York, New York, USA; Institute of Cancer Sciences, University of Glasgow, Glasgow, UK; Laboratory for Molecular Cancer Biology, VIB Center for Cancer Biology, VIB, Herestraat 49, 3000 Leuven, Belgium; Laboratory for Molecular Cancer Biology, Department of Oncology, KU Leuven, Herestraat 49, 3000 Leuven, Belgium; Department of Physiology, Yong Loo Lin School of Medicine, National University of Singapore (NUS), Singapore

**Author notes:** Correspondence should be addressed to Ernesto Guccione and Slim Mzoughi.

## Abstract

Targeting cancer stem cells (CSCs) is crucial for effective cancer treatment^1^. However, the molecular mechanisms underlying resistance to LGR5^+^ CSCs depletion in colorectal cancer (CRC)^2,3^ remain largely elusive. Here, we unveil the existence of a primitive cell state dubbed the oncofetal (OnF) state, which works in tandem with the LGR5^+^ stem cells (SCs) to fuel tumor evolution in CRC. OnF cells emerge early during intestinal tumorigenesis and exhibit features of lineage plasticity. Normally suppressed by the Retinoid X Receptor (RXR) in mature SCs, the OnF program is triggered by genetic deletion of the gatekeeper APC. We demonstrate that diminished RXR activity unlocks an epigenetic circuity governed by the cooperative action of YAP and AP1, leading to OnF reprogramming. This high-plasticity state is inherently resistant to conventional chemotherapies and its adoption by LGR5^+^ CSCs enables them to enter a drug-tolerant state. Furthermore, through phenotypic tracing and ablation experiments, we uncover a functional redundancy between the OnF and stem cell (SC) states and show that targeting both cellular states is essential for sustained tumor regression *in vivo*. Collectively, these findings establish a mechanistic foundation for developing effective combination therapies with enduring impact on CRC treatment.

## Main

CRC is currently the second leading cause of cancer-related mortality worldwide^4^. Treatment failure has traditionally been attributed to the malignant features of the LGR5^+^ cancer stem cells (CSCs) ^5–7^. However, recent evidence challenges this notion by highlighting a more prominent role of LGR5^-^ cells in the metastatic spread of CRC cells^8,9^. Furthermore, there is now unequivocal proof that targeting LGR5^+^ CSCs is insufficient to achieve a durable tumor regression^2,3^. Nevertheless, whether tumor evolution and adaptability in CRC are fueled by cellular plasticity, distinct cell populations, or a combination of both remains unclear.

### APC loss-of-function activates an OnF program and triggers lineage plasticity

To investigate whether new cell states emerge and persist during intestinal neoplasia, we generated isogenic mouse organoid models that mimic the clinical progression of human CRC (**Fig.1a and Extended Data Fig.1a**). By profiling the transcriptome of single cells from wild type intestinal organoids (WT), *Apc^KO^* (A) and *Apc^KO^::Kras^G12D^::Smad4^KO^::Trp53^KO^*(AKSP) tumoroids (**Extended Data Fig.1b**), we identified 24 distinct clusters (**Extended Data Fig.1c**). Intriguingly, cluster 22 (c22) contained a unique population of tumor-specific cells expressing markers of fetal intestinal progenitors^10^ (**Figs. 1b-c** **and Supplementary Table 1a)**. Henceforth, we refer to this population as oncofetal (OnF) cells and define a 91-gene signature (**Extended Data Fig.1d and Supplementary Table 1b**) to chart and track its dynamics during tumorigenesis. The changes in cellular composition along the CRC malignancy continuum were characterized by a progressive reduction in differentiated enterocytes, indicative of a differentiation block, and the emergence and persistence of OnF cells (**Fig.1d**). Reconstruction of cell state dynamics by RNA velocity analysis in premalignant (A) and cancerous (AKSP) tumoroids support the idea that OnF cells represent a more primitive state with the potential to give rise to all mature intestinal cell types including the LGR5^+^ SCs (**Fig.1e-f**). This finding is consistent with the emergence of a fetal-like cell state preceding the reappearance of LGR5^+^ cells in the regenerating epithelium^11^. While such retrogression occurs transiently in response to injury, our data indicate that APC loss-of-function (LoF) locks a subset of the premalignant LGR5^+^ SCs in an OnF state (**Fig.1d Extended Data** Fig. 1a, e**-f**).

**Figure. 1:**
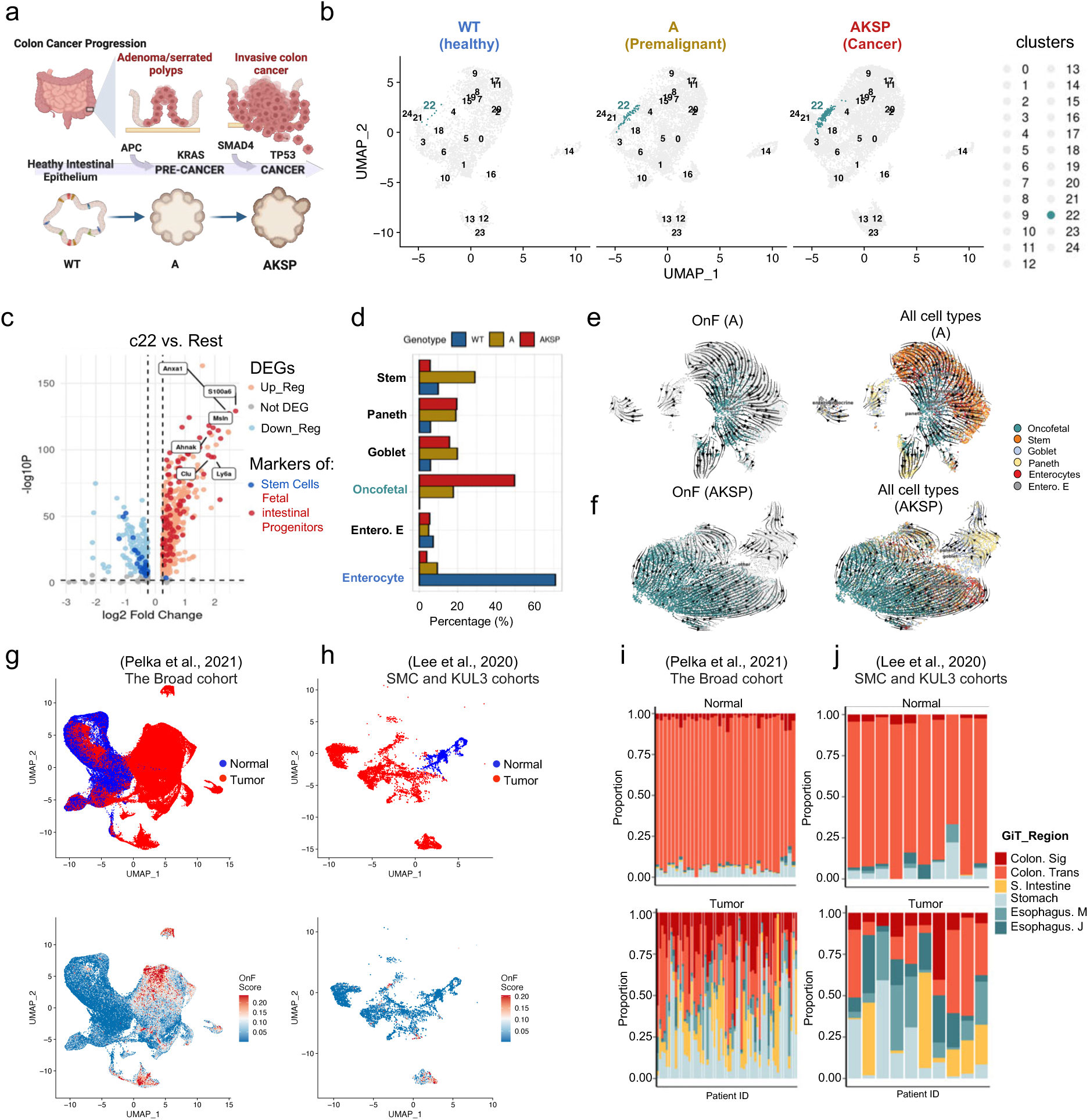
Identification of OnF cells, a neoplastic population with mixed lineage identity in CRC. **a,** schematics of the organoid models used to recapitulate the malignancy continuum of colorectal cancer. WT, Wild Type; A, *ApcKO;* AKSP*, ApcKO::Kras^G12D^::Smad4^KO^::Trp53^KO^.* **b,** Integrated UMAP of s.c RNA-seq data from the indicated organoid models; cells from the neoplastic cluster 22 (c22) are indicated in green. **c,** Volcano plot of significant differentially expressed genes (DEGs) in c22 vs. all other clusters. The dark blue and red colors correspond to markers of adult Stem Cell and Fetal progenitor genes, respectively. **d,** The percentage of various cell types and states in the indicated genotypes. Eentero.E, Enteroendocrine. **e, f,** Vector fields representing RNA velocity projected onto UMAPs of A **(e)** and AKSP **(f)** tumoroids. The color code indicates the spatial distribution of various cell states in each model. **g, h,** UMAP layouts of normal intestinal and tumor epithelial cells from the Broad **(g)** and SMC and KUL3 **(h)** CRC cohorts. In the upper panels, normal and tumor cells are indicated in blue and red, respectively. In the lower panel, cells were colored by expression of the Oncofetal signature (OnF score). **i, j,** Proportions of cells expressing specific signatures of the indicated gastrointestinal-tract tissues, in individual patients from the Broad **(i)** and SMC and KUL3 **(j)** cohorts. Sig, sigmoid, Trans, transverse; S, Small; M, Mucosa; J, junction.

Next, we defined a human OnF signature (**see methods and Supplementary Table 1c**) to investigate the emergence of this cellular state in patient-derived tumors. Analysis of single cell data from the SMC, KUL3 and Broad CRC cohorts^12,13^ confirmed that OnF cells are abundant in tumors while extremely rare in healthy intestinal tissues (**Fig. 1g-h** **and Extended Data** Fig. 1g). Moreover, we validated the tumor-specific enrichment of fetal-intestine genes in the TCGA/COAD dataset, which comprises 455 human colorectal adenocarcinomas and 41 normal colons (**Extended Data** Fig. 1h-i). Interestingly, comparing the transcriptome of CRC tumors with various developmental stages of the human gastrointestinal tract revealed striking similarities between tumors, fetal intestines, and adult stomach (**Extended Data Fig. 1j**). Overall, we noted a distinct transition of the neoplastic tissue from its posterior colonic identity towards more anterior regions of the gut tube (**Extended Data Fig.1k**). By analyzing single cell transcriptomic data from individual patients using lineage-specific signatures, we further confirmed that CRC tumors have acquired a metaplastic multiregional identity **(Fig.1i-j)**.

Activation of a fetal-like program and lineage infidelity have been independently reported in different regenerative contexts^11,14^. However, our data reveals that both processes co-occur in neoplasia, suggesting that they are related. Collectively, these findings indicate that APC LoF triggers a primitive cell state with a multiregional identity during the initial stages of intestinal tumorigenesis, priming tumor cells to adapt under selective pressure.

### Molecular determinants of the OnF state in CRC

Transcriptional programs that define various cellular states and identities are governed by transcription factors (TFs) networks that cooperate with epigenetic regulators to rewire the chromatin landscape. To investigate the molecular mechanisms underlying neoplastic cell states in CRC, we analyzed the transcriptome and chromatin accessibility landscapes along the malignancy continuum (**Extended Data** Fig. 2a). Transcriptomics data revealed that most gene expression changes coincided with depletion of the gatekeeper gene *Apc* at the premalignant stage (**Extended Data Fig. 2b**) and persisted throughout the genetic progression **(Fig. 2a, Extended Data** Fig. 2c **and Supplementary Table 2a)**. These transcriptional changes were consistent with the conspicuous reconfiguration of cell states identified by scRNA-seq (**Fig.1d)**, particularly highlighting the emergence of OnF cells (**Fig. 2b**).

**Figure. 2:**
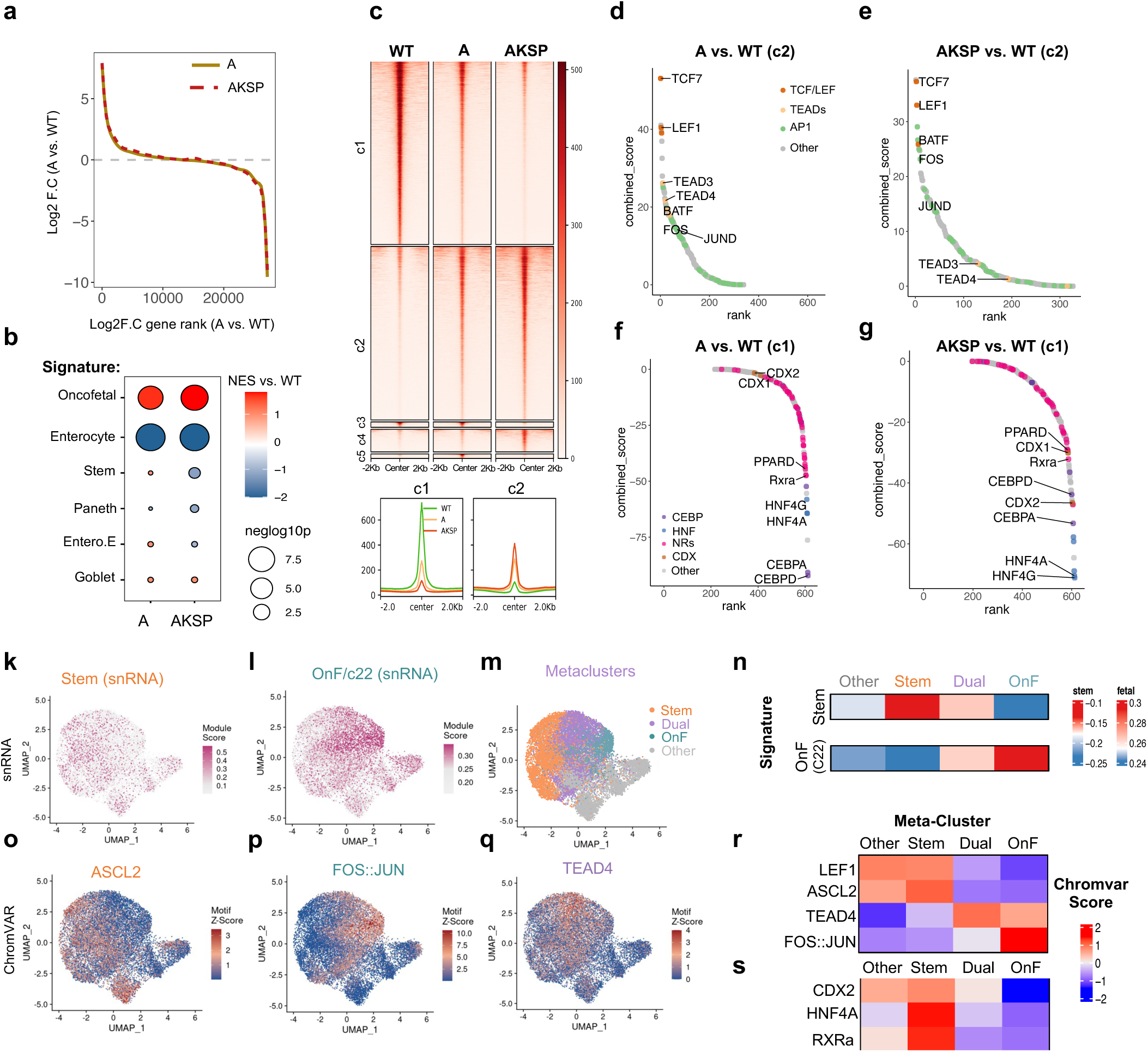
YAP and API drive OnF reprograming during intestinal neoplasia. **a,** Comparison of differential gene expression (log_2_ fold change) in AKSP vs. WT (dashed line) with A vs. WT as a reference (solid line). geom_smooth function from ggplot2 package is used to fit the curve to fold changes in reference comparison. **B,** Bubble plot depicting the enrichment of cell state-specific signatures in A vs. WT and AKSP vs. WT organoids using GSEA. Red and blue indicate positive and negative normalized enrichment scores (NES), respectively. Bubble size indicates the negative of log_10_ adjusted p-value. **c,** Heatmap of ATAC-seq signal in WT, A and AKSP organoids at differentially accessible regions (upper panel). Comparison of average ATAC-seq signal profile in the indicated organoid lines within a +/-2kb region around the peak center in clusters c1 and c2 (lower panel). **d, e, f, g**, Elbow plots depicting transcription factor (TF) activity in A vs. WT **(d, f)** and AKSP vs. WT **(e, g)**. TF combined binding score is defined as -log_10_(p-value) * log_2_(fold change) on the y-axis and ordered by TF rank on the x-axis. Differentially accessible regions from clusters c2 and c1 were used for analysis in panels (**d, e)** and (**f, g)**, respectively. **k, l,** UMAP embeddings of scATAC-seq cells from AKSP tumoroids. Enrichment of stem **(k)** and OnF/c22 **(l)** transcriptional signatures on snRNA-seq cells projected into the scATAC-seq manifold. **m,** Same UMAP from **(k, l)** colored by cell states as defined by the stem and OnF module scores. **n,** Enrichment of the stem (top) and OnF/c22 (bottom) module scores in the indicated meta-clusters. **o, p, q,** TF motif activity in scATAC-seq cells determined by chromVAR deviation scores for the indicated TFs. **r, s** TF motif activity in the indicated meta-clusters as defined by chromVAR scores (scores are scaled by row).

Similarly, changes in chromatin accessibility induced by APC LoF were largely maintained across the adenoma-adenocarcinoma sequence (**Extended Data Fig. 2d**). Unsupervised hierarchical clustering of ATAC-seq peaks revealed two primary patterns of chromatin accessibility (**Fig. 2c**). Cluster 1 (c1) consisted of genomic regions that exhibited reduced accessibility in the mutant tumoroids, while those in cluster 2 (c2) progressively gained accessibility (**Fig. 2c**). In contrast to events in clusters 3 - 5 (c3-c5), the persistent nature of these changes suggests they more likely reflect the emerging neoplastic cell states (**Fig. 2b**). To gain deeper insights into the functional significance of these changes, we performed TF footprinting analysis on both sets of ATAC-seq peaks, using TOBIAS^15^, and defined a “combined binding score” to assess TF activity across the malignancy continuum **(see methods)**. We found that the tumoroid-specific events in c2 were predominantly driven by three TF families (TCF/LEF, TEADs and AP-1) **(Fig.2d-e, Extended Data** Fig. 2e-h **and Supplementary Table 2b)**. The former operates under the canonical WNT signaling and, together with ASCL2, sustains the adult LGR5^+^ SC state^16^. TEADs, on the other hand, are the cognate DNA-binding partners of YAP, which has recently been implicated in the transient activation of a fetal-like program following injury^17,18^. Notably, re-activation of fetal genes is tightly associated with inflammatory responses^11,17^. Yet, despite the well-established role of AP-1 in regulating inflammation and evidence of its synergistic cooperation with YAP in other contexts^17,19,20^, the attractive possibility involving AP-1 in fetal regression has gone unexplored. Interestingly, HOMER motif analysis revealed a specific enrichment of both AP-1 and TEAD binding sites in the promoter region of OnF genes (**Extended Data** Fig. 2i). Together, these data suggest a cooperative view whereby AP-1 and YAP act in concert to establish the OnF state in intestinal tumors.

Conversely, genomic regions that became less accessible in tumoroids (c1) were enriched for footprints of the caudal-related homeobox (CDX) and hepatocyte nuclear factors (HNFs) family members **(Fig. 2f-g, Extended Data** Fig. 2j-m and **Supplementary Table 2c)** involved in establishing the caudal identity of intestinal cells and their maturation, respectively^21–23^. The diminished activity of these TFs is consistent with the anteriorization phenotype (**Fig. 1i-j** **and Extended Data** Fig. 2j-k) and the retrogression to a more primitive state in tumors **(Fig. 1d, g-h)**. Intriguingly, we also noted a significant reduction in the activity of several ligand-regulated nuclear receptors, including PPAR, RXR, LXR, VDR and FXR, all of which require dimerization with RXR to become functional **(Fig. 2f-g, Extended Data** Fig. 2l-m and **Supplementary Table 2c).**

To achieve a more comprehensive understanding of the molecular mechanisms underlying neoplastic states at the single cell level, we performed single cell Multiome (scATAC+scRNA-seq) analysis on CRC tumoroids (AKSP). We first projected scATAC-seq cells into low-dimensional subspaces and identified 16 clusters (**Extended Data** Fig. 2n). We then overlayed the OnF (c22) and stem cell transcriptional signatures (from scRNA-Seq data) onto the UMAP of chromatin accessibly to define spatial distribution of the various cell states. This analysis allowed us to identify distinct meta-clusters representative of the OnF and stem cell states (**Fig. 2k-l**) and unveiled the existence of an intermediate cell population in a dual (OnF/Stem) cell state (**Extended Data** Fig. 2o **and** **Fig. 2m-n**). This hybrid profile bridging the OnF and stem cell compartments suggests that neoplastic cells exist in a metastable state, in absence of selective pressure. To further investigate TFs that govern these dynamic cell states, we used the chromVAR package^24^ to map TF motif activity across the various topological domains. This analysis confirmed that ASCL2 binding sites were primarily accessible in SCs (**Fig. 2o, r**). Interestingly, while JUN::FOS motif activity was exclusive to OnF cells (**Fig. 2p, r**), TEAD4 had a less selective profile and was rather characteristic of the transitional state (**Fig. 2q-r**). These findings support an adaptive bet-hedging model, in which YAP-signaling plays the role of epigenetic switch facilitating the transition to an OnF state, further reinforced by AP-1 activation. Noticeably, the progressively reduced accessibility of CDX2 and HNF4 binding sites in cells with an active OnF program (**Fig. 2s** **and Extended Data** Fig. 2p-q) was consistent with the gradual loss of the mature intestinal identity. However, we were again intrigued by the motif activity pattern of RXR, which was instead equally silenced in OnF and dual state cells (**Fig. 2s** **and Extended Data** Fig. 2r), suggesting an early regulatory role in OnF regression.

### RXR is a gatekeeper of OnF reprogramming in CRC

To investigate whether the diminished RXR activity following APC LoF contributes to the onset of the OnF state, we conducted a series of comparative analyses between RXR inhibition and APC depletion. Treatment with the RXR antagonist HX531, hereafter referred to as RXRi, altered the budding structure of WT organoids (**Fig. 3a**) and imposed a spheroid morphology (**Fig. 3b**) reminiscent of both fetal organoids^10^ and mutant tumoroids (**Fig. 3c**). Transcriptomic analysis revealed a remarkable resemblance between RXRi organoids and both mouse tumoroids (**Fig. 3d**) and human CRC (**Fig. 3e**), while WT organoids were more similar to healthy colonic tissues (**Fig. 3e**). In addition, transcriptional signatures of fetal intestinal progenitors, gastric, and esophageal metaplasia were significantly enriched in RXRi organoids (**Extended Data Fig. 3a-c**). Specifically, RXR inhibition resulted in a global shutdown of all mature cell markers, including stem cells, and induced a more prominent increase in fetal genes (**Fig. 3f**). This is in contrast to APC depletion, which, in addition to causing a block in differentiation and activation of the OnF program, induced an expansion of the mature LGR5^+^ SCs (**Fig. 3f** **and** **1d**).

**Figure. 3:**
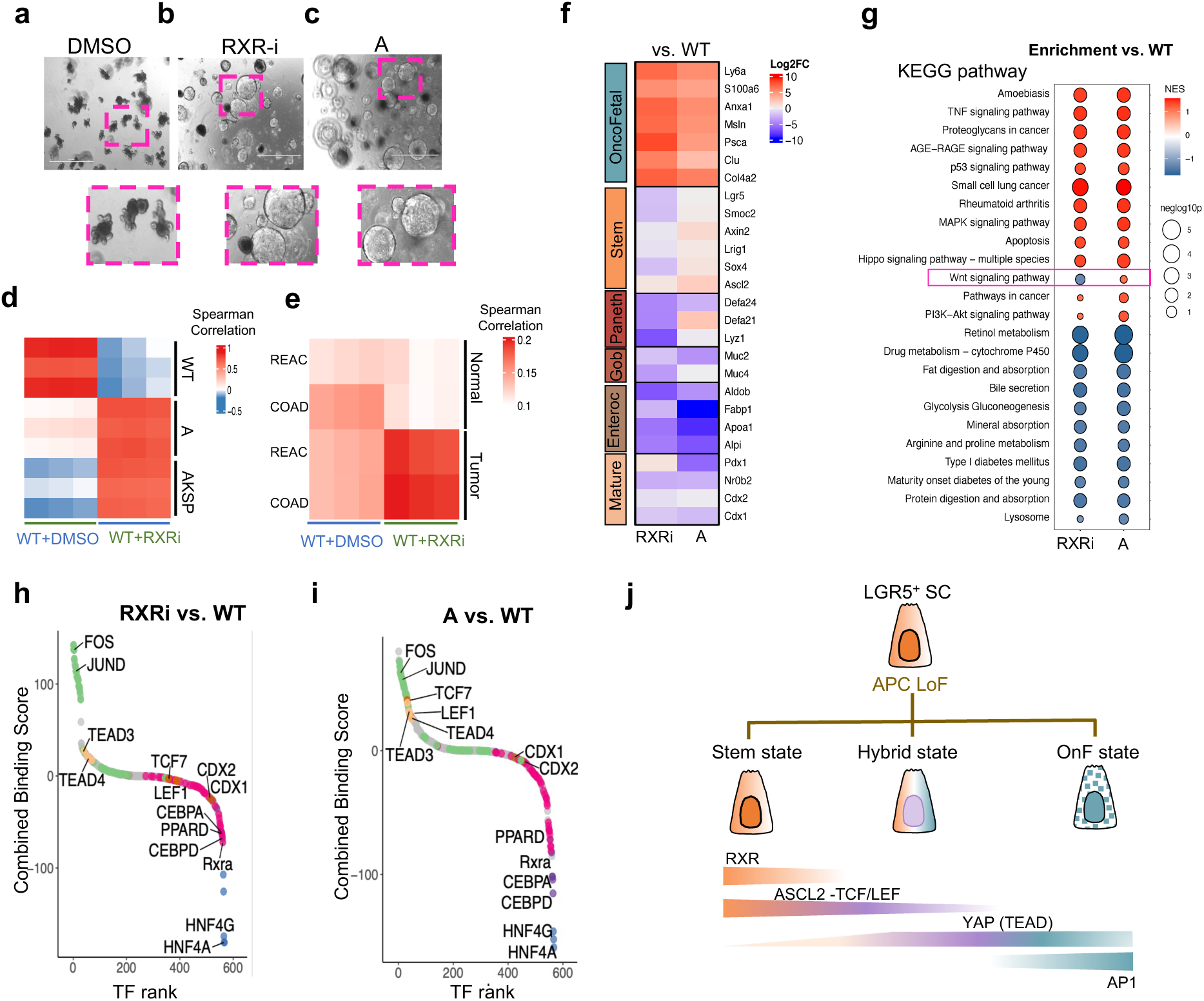
RXR opposes oncofetal reprogramming by counteracting YAP/AP1 activity. **a, b, c,** Representative images of WT organoids treated with DMSO **(a)** or the RXR antagonist HX531 (RXRi) **(b)** and A tumoroids **(c).** Lower panels are a close-up of the dashed line areas highlighting the morphological differences and similarities between the different conditions. **d,** Spearman correlation of the top 1000 highly variable genes in DMSO, RXRi treated organoids, mouse healthy organoids (WT) and mutant tumoroids (A and AKSP) **(d)** or top 2000 highly variable genes in DMSO, RXRi treated organoids, human healthy colons and tumors from the COAD and REAC TCGA datasets **(d)** or human healthy colons and tumors from the COAD and REAC TCGA datasets **(e). f,** Heatmap of differentially expressed marker genes of the indicated mature intestinal and oncofetal cell states in RXRi vs. WT and A vs. WT. **g,** GSEA of KEGG pathway enrichment in RNA-seq data from RXRi vs. WT and A vs. WT. Bubble color and size indicate normalized enrichment score (NES) and -log_10_(p-value), respectively. **h,i,** Elbow plots depicting transcription factor (TF) activity in RXRi vs. WT **(h)** and A vs. WT **(i).** TF combined binding score is defined as -logw(p-value) * log2(fold change) on the y-axis and ordered by TF rank on the x-axis. **j,** Proposed model illustrating the establishment of a spectrum of neoplastic cell states through opposing gradients of RXRa and YAP/AP1 activities following APC Loss-of-Function (LoF). The dotted pattern in the OnF state is to reflect the mixed lineage identity.

Functional analysis of the RNA-seq data revealed that RXR inhibition recapitulated most transcriptional changes induced by APC LoF with the main difference being a divergent regulation of the WNT signaling pathway **(Fig. 3g, Extended Data** Fig. 3d-e and **Supplementary Table 3a)**. This may explain the more pervasive regression to an OnF-like state observed in RXRi organoids (**Fig. 3f**). To gain deeper understanding of how RXRa inhibition favors a complete reversion to an OnF-like state, we investigated its effects on global TF dynamics. First, ATAC-seq data unveiled significant similarities in chromatin accessibility landscapes between *Apc* mutant tumoroids and RXRi organoids (Extended Data Fig. 3f-h). Second, TOBIAS footprinting showed that RXR inhibition recapitulated most TF activity changes induced by APC LoF (**Fig. 3h-i** **and Supplementary table 3b)**, including reduced activity of RXRa itself, CDX2, and HNF4 (**Extended Data Fig. 3k-m**) and hyperactivation of the OnF-specific TFs AP-1 and TEADs (**Extended Data Fig. 3n-o**). However, unlike APC depletion, RXR blockade did not activate WNT-related TFs (**Fig. 3h-i** **and Extended Data** Fig. 3p). HOMER analysis confirmed that regions with decreased accessibility in RXRi organoids were enriched for the TCF/LEF TF DNA binding motif (**Extended Data Fig3. q and** **Supplementary Table 3c)**. Together with the scATAC+scRNA-seq Multiome data, these results support a model wherein the fate of *Apc* mutant SCs is determined by the level of RXRa activity (**Fig. 3j**).

Overall, our findings unveil an antagonistic relationship between RXRa and YAP/AP-1 and posit the former as a gatekeeper of the OnF state. Furthermore, we show that varying gradients of OnF and SC drivers generate a continuum of neoplastic cellular states promoting phenotypic heterogeneity (**Fig. 3j**).

### Visualization and targeting of the OnF state in CRC

It is noteworthy that despite enabling the inference of cellular states^25^, transcriptional signatures offer only a static snapshot. To comprehensively understand the temporal dynamics of these states, a well-defined and measurable tool is needed. We have recently developed a new strategy to genetically trace cell fate transitions within a heterogenous tissue^26^. To this end, we leveraged our knowledge of the molecular underpinnings of the OnF program to construct a synthetic locus control region (sLCR) that contains specific cis-regulatory elements (CREs) reflecting the transcriptional output and activity of OnF state-associated TFs (**Extended Data Fig. 4a-b** and **Supplementary Table 4a)**. Then, we fused this genetic tracing cassette to an enhanced Green Fluorescent Protein (eGFP) (**Fig. 4a**) to allow visualization and tracking of this cellular state. Flow cytometry analysis of WT organoids expressing this phenotypic reporter confirmed that only a small fraction of cells exhibited low to moderate levels of GFP expression (**Extended Data** Fig. 4c). In contrast, WT organoids treated with RXRi and tumor-derived VAKSP (Villin-Cre AKSP) organoids displayed a notable shift in reporter activity, evidenced by higher proportion of GFP^+^ cells and increased Mean Fluorescence Intensity (MFI) (**Extended Data Fig. 4c-d**). Moreover, transcriptomic analysis of sorted GFP^high^ and GFP^negative^ cells from VAKPS tumoroids confirmed the enrichment of the OnF signature, (c22)-specific genes, and markers of lineage plasticity in the former (**Fig. 4b** **and Extended Data** Fig. 4e **and Supplementary Table 4b**). These data strongly support the validity and specificity of our phenotypic reporter as an effective tool for tracing OnF cells.

**Figure. 4:**
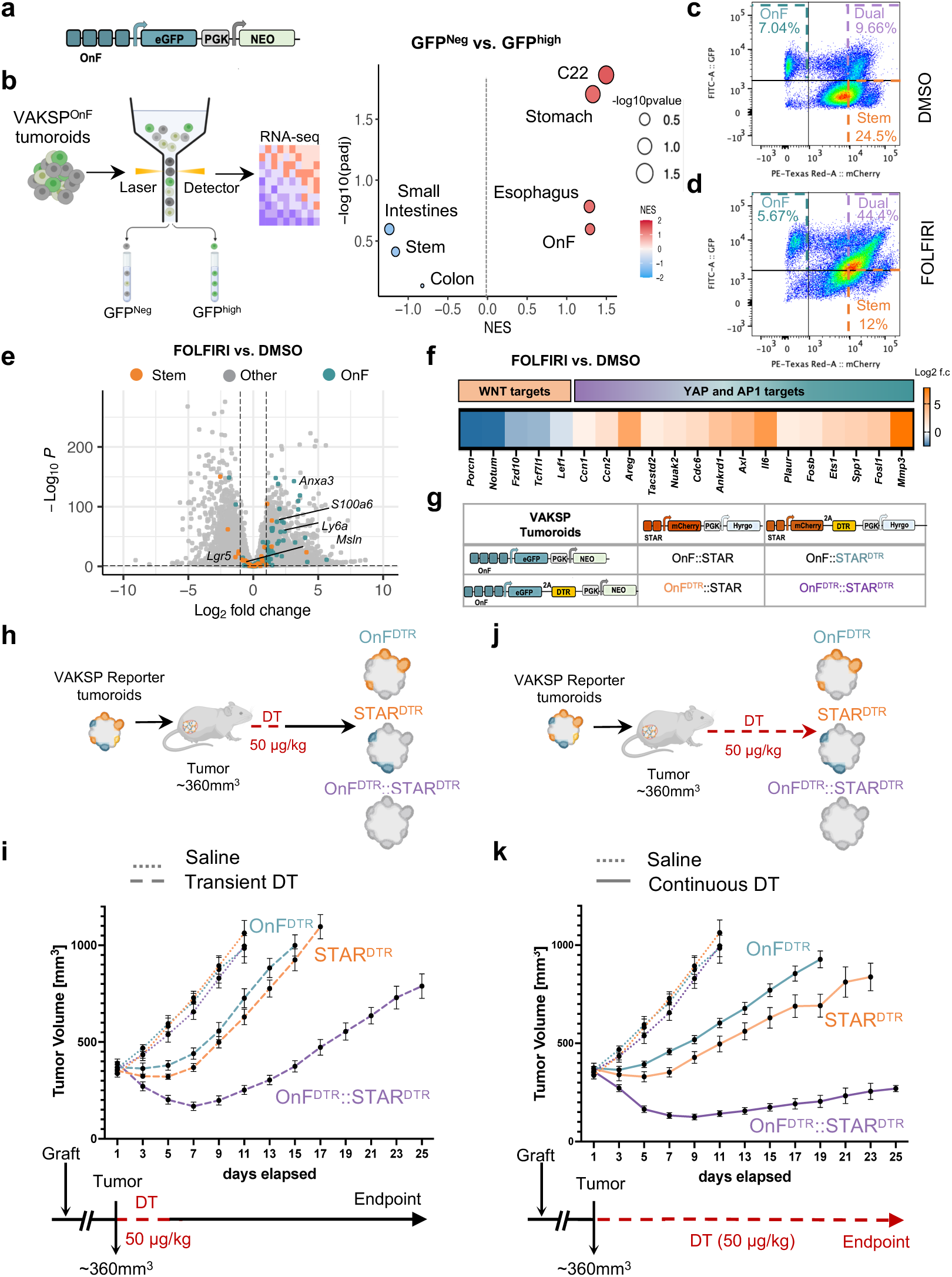
A functional interplay between OnF and stem cells fuels phenotypic plasticity in CRC. **a,** Schematic of the oncofetal (OnF) phenotypic reporter structure**. b,** Schematic representation of the experimental flow (left) and enrichment analysis of the indicated gene-set signatures in GFPhigh compared to GFPNeg cells sorted from VAKSP (Villin-Cre AKSP) tumoroids expressing the OnF reporter (VAKSPOnF) (right). Red and blue indicate positive and negative normalized enrichment scores (NES), respectively. Bubble size indicates the negative log_10_ p-value. **c, d,** Representative pseudocolor plots of flow cytometry analysis of VAKSP tumoroids co-expressing the oncofetal (OnF-GFP, Y axis) and stem (STAR-mCherry, X axis) cell phenotypic reporters. VAKSP ^OnF/STAR^ tumoroids were treated with DMSO **(c)** or FOLFIRI **(d)** for 3 days before analysis. **e,** Volcano plot of differentially expressed genes (DEGs) in VAKSP tumoroids following 3 days of FOLFIRI treatment compared to DMSO control. Significant DEGs are located on both sides of the dashed lines (log2FC is 1 and p-value <0.01). OnF and Stem cell markers are indicated in green and orange, respectively. **f,** Heatmap reporting log_2_ fold change in WNT (left) and YAP/AP1 (right) target gene expression. **g,** A summary of the VAKSP tumoroid models generated for downstream functional studies. **h, j,** Schematic representation of the experimental strategy to genetically target the OnF state, stem state, or both, relevant to panels **(i)** and **(k)**, respectively. **i, k,** Growth rate of subcutaneous VAKSP tumors expressing the indicated reporter combinations, in response to Diphtheria toxin (DT) treatment. Mean tumor volume ± s.e.m are shown. Dotted lines indicate saline treatment (all models, n=6 tumors). **i,** Dashed lines indicate transient (5 days; 14 doses) DT treatment; OnFDTR (n=11); STARDTR (n=9); OnFDTR /STARDTR (n=10). **k,** Solid lines indicate continuous DT treatment; OnFDTR (n=12); STARDTR (n=11); OnFDTR/STARDTR (n=9). Bottom panels show the dosing schedule (every other day) of DT -duration of treatment is indicated by a dashed red line.

Next, to investigate their interplay with LGR5^+^ SCs, we replaced the OnF sLCR cassette with the STAR minigene^27^ driving mCherry expression (**Extended Data Fig. 4f**) and generated a tumoroid line that expressed both reporters (VAKSP^OnF/STAR^). This strategy allowed us to visualize the continuum of neoplastic cell states including the (OnF/SC) hybrid cells (**Extended Data Fig. 4g-h**). Moreover, we captured dynamic shifts between SCs (STAR^high^) and the dual (OnF^+^/STAR^high^) population at steady state conditions (**Extended Data** Fig. 4h-j), suggesting that phenotypic plasticity is an inherent feature of LGR5^+^ SCs. To unravel the functional relevance of these transitions, we treated VAKPS^OnF/STAR^ tumoroids with FOLFIRI, a commonly used chemotherapy combination in clinical practice. Analysis of cell state dynamics using flow cytometry revealed a substantial transition of SCs to a hybrid state, while OnF cells remained largely unaffected by therapeutic pressure (**Fig. 4c-d**). These data are consistent with the observed signs of lineage plasticity (**Extended Data** Fig. 4k), upregulation of key OnF markers (**Fig. 4e** **and Supplementary Table 4c)** and YAP/AP-1 target genes (**Fig. 4f** **and Supplementary Table 4c)** in response to FOLFIRI. They are also in agreement with the reduced expression of WNT-related genes (**Fig. 4f**). These observations indicate that the OnF state, exhibiting a multiregional identity, is inherently resistant to chemotherapies, and that activation of the associated program in SCs allows them to escape treatment.

Notably, recent evidence has shown that ablation of the LGR5^+^ SCs altogether is insufficient to achieve a durable regression of CRC^2,3^, which suggests that targeting their phenotypic plasticity alone may not be enough to overcome resistance. Our data (**Fig. 4c-d**) imply that in such scenario, the pre-existing OnF cells may play a crucial role in sustaining tumor growth. To test this hypothesis, we developed a Diphtheria Toxin Receptor (DTR)-expressing version of the OnF and SC tracing cassettes. We then generated VAKSP tumoroids that express various combinations of these reporters to allow selective attrition of either or both cellular states (**Fig. 4g**). In absence of DTR expression, Diphtheria Toxin (DT) treatment did not affect cellular composition nor viability of VAKSP^OnF/STAR^ tumoroids (**Extended Data Fig. 4l, p**). However, the respective target populations were efficiently ablated in the DTR^+^ models (**Extended Data Fig. 4m-o** **and q-s)**. Importantly, this cell state-specific strategy enabled simultaneous ablation of hybrid cells (**Extended Data Fig. 4m-o**), effectively targeting phenotypic plasticity.

To investigate the therapeutic potential of targeting the OnF state in CRC, we transplanted the different VAKPS reporter lines (**Fig. 4g**) into the flanks of immune-compromised mice. We first allowed tumors to reach a relatively large volume (350-400mm^3^) before we started DT administration every other day for 5 days (**Fig. 4h-i**). Consistent with previous work^2,3^, selective ablation of cells with an active SC program led to tumor stasis in the VAKSP^STAR-DTR^ model, followed by a prompt regrowth upon DT cessation (**Fig. 4i**). Similarly, attrition of OnF^+^ cells halted tumor growth transiently but was not sufficient to cause tumor regression (**Fig. 4i**). In contrast, co-targeting both cell states caused tumor shrinkage and delayed resurgence significantly (**Fig. 4i**). Intriguingly, continuous depletion (**Fig. 4j-k**) of OnF^+^ or STAR^+^ cells only failed to sustain tumor stasis indefinitely (**Fig. 4k**), suggesting that either population is capable of fueling tumor growth in absence of the other. However, we noted a prominent tumor regression that was maintained for as long as DT treatment continued in the VAKSP^OnF-DTR/STAR-DTR^ model (**Fig. 4k**). Collectively, these data unveil a functional redundancy between OnF and Stem cells and underline the therapeutic potential of a combinatorial approach targeting molecular programs that govern both cellular states in CRC.

## Discussion

Following the discovery of LGR5^+^ intestinal SCs^28,29^, their role as the driving force behind tumor growth^30–32^ has spurred the design of CSCs therapies^33,34^. Such strategy assumes that CRC displays the same rigid cellular hierarchy of the homeostatic intestinal epithelium. Our studies challenge this assumption and identify OnF cells as a distinct neoplastic state. Driven by YAP and AP-1 activity, these cells display features of lineage plasticity and are inherently resistant to chemotherapies. Moreover, activation of this program allows LGR5^+^ CSCs to enter a drug tolerant state.

Interestingly, mixed lineage states resembling OnF cells have been recently observed in pancreatic^35,36^, lung^37–39^ and prostate cancer^40^ at various stages of tumor evolution. Therefore, we propose that the early onset of a highly plastic OnF state represents a universal mechanism of cellular plasticity in cancer. This assumption is further supported by the role of the OnF drivers, YAP and AP-1, as orchestrators of an aberrant pan-cancer enhancerome^19,20,41–43^. It is worth noting that although the inflammatory circuits at play may be context-dependent, the activation of regenerative and inflammatory pathways emerge as a recurring mechanism in promoting epigenetic plasticity^36,40,44^.

Importantly, our studies provide compelling evidence that RXR acts as a master regulator of these pathways and serves as a gatekeeper of OnF regression during intestinal tumorigenesis. This suggests that the protective effects of statins in hyperlipidemia patients^45,46^ may be attributed to the early blockade of the OnF program through activation of the RXR::PPAR complex. In fully developed CRC, however, we demonstrate that targeting the OnF state alone has marginal effects on tumor growth. These data complement previous studies indicating the limited potential of CSC-targeting therapies^2,3^ and uncover a functional redundancy between these two cellular states. Furthermore, we provide strong evidence that targeting both the SC and OnF programs induces a durable regression of CRC, effectively locking tumors in a minimal residual disease state (**Fig. 4k**). These findings establish a foundation for targeted drug-repurposing screens to identify effective combination therapies. In addition, our data from organoid models suggest that OnF reprogramming in the tumor likely precedes that of its ecosystem^47^, although the mechanisms underlying such crosstalk remain to be determined.

## Reporting Summary

Further information on research design is available in the Nature Research Reporting Summary linked to this paper.

### Data Availability

All data needed to evaluate the conclusions in this study are present in the paper and/or its Supplementary Materials. RNA-seq, ATAC-seq, scRNA-seq, scATAC-seq and CUT&RUN data supporting the findings of this study have been deposited into the National Center for Biotechnology Information (NCBI) Gene Expression Omnibus under series accession GSE235205.

## Acknowledgments

Research reported in this publication was supported in part by ISMMS seed fund to EG. The authors gratefully acknowledge use of the services and facilities of the Tisch Cancer Institute supported by the NCI Cancer Center Support Grant (P30 CA196521).

This work was supported in part by the Bioinformatics for Next Generation Sequencing (BiNGS) shared resource facility within the Tisch Cancer Institute at the Icahn School of Medicine at Mount Sinai, which is partially supported by NIH grant P30CA196521. This work was also supported in part through the computational resources and staff expertise provided by Scientific Computing at the Icahn School of Medicine at Mount Sinai and supported by the Clinical and Translational Science Awards (CTSA) grant UL1TR004419 from the National Center for Advancing Translational Sciences. Research reported in this paper was supported by the Office of Research Infrastructure of the National Institutes of Health under award number S10OD026880.

This research was supported by an International Accelerator Award, ACRCelerate, jointly funded by Cancer Research UK (A26825 and A28223), FC AECC (GEACC18004TAB), and AIRC (22795).

## Author contributions

SM and EG conceptualized the project and designed experiments. SM, MS and FDT performed the experiments and acquired the data. SM and KM prepared ATAC-seq and RNA-seq libraries, respectively. DD, XW, GU, DH, AT, BG performed computational and omics analysis with input from SM and EG. YD, CC and GG designed the oncofetal reporter with input from SM. ML and OJS generated the VAKSP tumoroids. ML and NB generated the remaining genetic organoids models. DLO provided guidance for flow cytometry data collection and acquisition. All authors provided scientific input. SM and EG wrote the manuscript, with inputs from all authors, and supervised research.

## Competing Interests

The Guccione laboratory received research funds from AZ and Prelude Therapeutics (for unrelated projects), EG is a cofounder and shareholder of Immunoa Pte.Ltd and cofounder shareholder, consultant, and advisory board member of Prometeo Therapeutics.

## Methods

### Mouse strains

Used to derive the *Lgr5-2A-CreERT2* organoids: All animal experiments underwent approval from the Institutional Animal Care and Use Committee. *Lgr5-2A-CreERT2* (Leushacke et al., 2017) mice were generated through homologous recombination in embryonic stem cells, specifically targeting the insertion of the *2A-CreERT2* cassette at the stop codon of the *Lgr5-ORF*. To obtain the experimental mice, *Lgr5-2A-CreERT2* mice were crossbred with *Rosa26-tdTomato (Ai14)* (Madisen et al., 2010), *LSL-KrasG12D* (Johnson et al., 2001), *Apc-loxP flanked* (Shibata et al., 1997), *Trp53-loxP flanked* (Jonkers et al., 2001) and *Smad4-loxP flanked* (Yang et al., 2002) mice, which were acquired from the Jackson Laboratory. All mice were bred onto the C57Black6/J background. The animals were subjected to a 12-hour light-dark cycle and were maintained at a room temperature of 21 ± 1 °C with a humidity level of 55–70%. To ensure pathogen-free conditions, all mice were bred and housed in ventilated cages. For all experiments, mice aged a minimum of 8 weeks were selected, and experimental groups were randomly assigned sex-matched littermates.

Used to derive the *villin-Cre^ER^* tumoroids: The experiment was performed according to UK Home Office regulations (Project License 70/8646), adhered to ARRIVE guidelines and was reviewed by local animal welfare and an ethical review committee at the University of Glasgow. Intracolonic induction in a 12-week-old male mouse *villin*Cre^ER^ *Apc*^fl/+^ *Kras*^G12D/+^ *Trp53*^fl/fl^ *Smad4*^fl/fl^ on a C57BL/6 background was performed under general anesthesia. A single 70µl 100uM dose of 4-hydroxy tamoxifen (H7904-5MG from Sigma) was injected into the colonic sub-mucosa via a colonoscope (Karl Storz TELE PACK VET X LED endoscopic video unit). At clinical endpoint (weight loss with or without the presentation of hunching) colonic tumor tissue was collected, and organoid cell lines were generated as previously described (Jackstadt et al., 2019).

### Organoid derivation and culture

Organoid lines were generated from small intestinal crypts, isolated from the respective mouse models, as previously described^29^. Briefly, the initial part of the small intestines was opened longitudinally and washed with phosphate-buffered saline (PBS). After removing the villi using a cold glass slide, the tissue was cut into small fragments (∼2mm in length) and incubated in 2.5 mM EDTA solution while shaking (4°C for 30 minutes). After removing the supernatant, the fragments were resuspended in advanced DMEM/F12 with 0.1% BSA and vigorously shaken. The suspension was then passed through a 70 μm strainer to collect the first fraction. The remaining tissue fragments were collected from the strainer and subjected to a second round of vigorous shaking before being passed through another 70 μm strainer. This process was repeated until 4 fractions were collected. Each of these was then centrifuged at 300g for 5 minutes at 4°C and the respective pellets were re-suspended in 7mg/ml Matrigel (Corning cat#35234) and plated into 24 well plates. Matrigel was allowed to polymerize at 37 °C for 10-15 minutes before mouse IntestiCult Organoid Growth Medium (STEMCELL Technologies cat#06005), containing Y-27632 (10 μM), was added. Organoids were passaged every 3-4 days by dissolving the Matrigel in Gentle Cell Dissociation Reagent (GCDR - STEMCELL Technologies cat#100-0485). Organoids were broken down to smaller sizes by trituration in GCDR before washing with basal media (advanced DMEM/F12, N2/B27 supplements, N-acetylcysteine, 1mM HEPES and 100 μg/ml Penicillin-Streptomycin). Cell pellets were then resuspended in Matrigel.

### Isogenic tumoroid line generation and treatment

To generate *Apc^KO^* tumoroids, *Lgr5*-2a-CreERT2::*Apc^flox/flox^* healthy organoids were treated with 4-hydroxytamoxifen (4-OHT, 500nM) (Sigma cat#SML1666-1ML) for 24hrs - intestinal stem cells with successful Cre-recombination activated tdTomato expression and appeared red under a fluorescence microscope. These tumoroids were then cultured in BEN selection media (basal medium with mouse recombinant EGF (50 ng/ml) and Noggin (100 ng/ml) (PeproTech) without R-spondin) for approximately 3 passages. Wild type (WT) organoids cultured in same medium were used as a control for selection. To generate the *Apc^KO^::Kras^G12D^::Smad4^KO^::Trp53^KO^*(AKSP) tumoroid line, *Lgr5-2a CreERT2::Apc^flox/flox^::Lox-Stop-Lox-Kras^G12D^::Smad4^flox/flox^*organoids were first treated with 4-OHT. The resultant AKS tumoroid line was selected in basal medium only, then edited by CRISPR–Cas9 to knock-out the *Trp53* gene. Following transfection with the PX458 (pSpCas9 (BB)-2A-GFP) vector expressing the targeting double-stranded gRNAs, they underwent a second round of selection in medium containing 10 μM Nutilin-3 (Sigma). Once established, these lines were maintained in complete mouse IntestiCult Organoid Growth Medium (STEMCELL Technologies).

Retinoid X Receptor (RXR) blockade experiments: WT organoids were initially replated in 1.25 μM of the RXR antagonist HX531 and then maintained in half that dose following the first passage. FOLFIRI treatment experiments: Tumoroids were treated with a 5uM mix of (5FU and Irinotecan; 1:1) (Selleck Chemicals) for 3 days before collection for downstream analyses.

Diphtheria Toxin (DT) treatment: VAKPS tumoroids expressing the indicated combination of phenotypic reporters were treated with 100ng/ml of DT or an equivalent volume of demineralized water (demi-water) for 3 days before collecting for downstream analyses.

### Phenotypic reporter design and generation

The OnF sLCR was designed using the logical design of synthetic cis-regulatory DNA (LSD) method, as described in (https://www.biorxiv.org/content/10.1101/2022.11.04.515171v1; https://gitlab.com/gargiulo_lab/s <u>LCR_selection_framework</u>), with custom deviations from recommended use of the pipeline. First, we focused on three biomarkers only: S100a6, Anxa1, Ly6a, as these were more specific to the OnF population. We retrieved the genomic coordinates of their boundaries (S100a6: chr3:90,612,893-90,614,414; Anxa1: chr19:20,373,433-203,906,71; Ly6a: chr15:749,948,76-749,980,31) from mm10 genome reference (UCSC genome browser RefSeq table). Second, we curated a list of transcriptional regulators based on differential gene expression between candidate OnF and Stem populations, Ingenuity Pathway analysis and Gene Ontology. Finally, we queried their respective known transcription factor binding sites (TFBS) and associated PWM weight matrices (PWMs). Together, this information allowed LSD to generate a pool of potential cis-regulatory elements (CREs) with a fixed length within user-defined regulatory landscapes (determined by CTCF sites surrounding each signature gene) [default is a 150–base pair (bp) window sliding with a 50-bp step]. Next, LSD assigned TFBSs to the CRE pool using FIMO (default --output-pthresh 1e-4 --no-qvalue) and created a matrix of putative CREs × TFBS. Finally, LSD selected the minimal number of CREs representing the complete set of TFBS by sorting and selecting the best CRE by the sum of the affinity score [−log10(P value)] and TFBS diversity (number of different TFBS). The first CRE was prioritized based on 5′ CAGE data (ENCODE) to increase the chances of successful transcriptional firing. From the rank-ordered list, we selected the top 7 CREs, and synthesized these in that order, using the same strand for “plus” and reverse and complement for “minus”.

The oncofetal sLCR was synthetized and cloned into a 3^rd^ generation pLV lentiviral backbone at VectorBuilder. To generate the STAR-mCherry reporter, eGFP was substituted by mCherry (VectorBuilder) and the OnF sLCR was replaced with the STAR minigene (Addgene #136255) using BamHI restriction enzyme and Gibson cloning.

### Virus production and Transductions

Lentiviral packaging plasmids (pCMV-D8.9 and pCMV-VSVG) and reporter plasmids were transfected into HEK-293T cells using Lipofectamine 3000 (ThermoFisher cat#L3000001). Virus-containing supernatant was harvested at 48 and 72 hours after transfection and viral particles were concentrated with Lenti-X Concentrator according to the manufacturer’s instructions (Takara Bio cat#631232). Lenti-X GoStix Plus were then used to determine virus titer (Takara Bio cat#631280). For lentiviral infection, organoid/tumoroid lines were dissociated and lentivirus was added along with TransDux™ MAX Lentivirus Transduction Reagent (SBI cat#LV860A-1). These mixtures were then centrifuged at 600g; 32°C for 15 (organoids) and 60 (tumoroids) minutes and incubated in a humidified 37°C incubator for 6 hours. Cells were then washed with basal medium (500g, 4°C) and replated in Matrigel in mouse IntestiCult Organoid Growth Medium (STEMCELL Technologies) supplemented with Y-27632 (10 μM). 3-4 days post infection, the different lines and non-infected controls were selected using the relevant antibiotics. After the initial selection, the generated reporter lines were maintained in half dose antibiotics, which were removed from the culture medium before using for experiments.

### Flow cytometry analysis

Matrigel drops containing organoids/tumoroids were collected in cold basal media. Following centrifugation (500G for 5 mins at 4°C), cells were incubated in TrypLE at 37°C and then mechanically dissociated by pipetting. After washing with 5 mL of cold basal medium (500G for 5 minutes at 4°C), samples were resuspended in 1mL of FACS buffer (PBS with 2% FBS and 1mM EDTA) containing a live/dead dye diluted 1:2000 (Live/Dead Fixable Aqua, Invitrogen #L34957) and incubated for 10 min at RT. In some experiments, cells were resuspended in 1mL of PBS and live/dead dye used at 1:500 (Zombie NIR, BioLegend #423106). Cells were washed with 5mL of FACS buffer, passed through a 40uM cell strainer into a new 15mL conical tube, and centrifugated (500G for 5 minutes at 4°C). Pellets were resuspended in 200uL of FACS buffer, transferred to 5mL polystyrene FACS tube, and immediately acquired on a BD LSRFortessa Cell Analyzer (BD Bioscences). For sorting experiments, an additional filtering step (35μm) was added. Cells were sorted on a CytoFLEX SRT (Beckman Coulter). In all experiments, unstained cells and FMO controls were included to determine basal levels of fluorescence intensity. Data were analyzed using FlowJo software (version 10.7.1, Tree Star, Ashland, OR, USA).

### Subcutaneous transplantation

VAKSP tumoroids expressing various combination of the phenotypic reporters were dissociated into small clusters of cells and resuspended in a Matrigel: Basal media (1:1) admixture. The equivalent of 5x10^5^ cells was injected subcutaneously in both flanks of 7 to 8-week-old female NSG mice (Strain #:005557, Jackson Laboratory). Tumor dimensions were measured using calipers and volume was calculated as 0.5x(Length x width^2^). DT (Sigma, # 0564) was administered via Intra-peritoneal (i.p.) injection at a dose of 50 μg/kg every other day for the duration indicated in each experimental setup. Mice were closely monitored by the authors, facility technicians and by a veterinary scientist responsible for animal welfare. They were euthanized when they reached a humane endpoint as determined by the Institutional Animal Care and Use Committee (IACUC). Some of these criteria include clinical signs of persistent distress or pain, significant loss of body weight (> 20%), tumor size exceeding 1000mm^3^, or when tumors ulcerated.

### RNA-seq and library preparation

Organoids/tumoroids were grown in complete medium (mouse IntestiCult Organoid Growth Medium-STEMCELL Technologies) for 3-4 days before they were collected directly in TRIzol reagent (Invitrogen cat#15596026). Before collection, treatments were added as indicated in figure legends and the appropriate methods sections. RNA was extracted using the PureLink RNA Mini Kit (Ambion cat#1283-018A) and quality was assessed with a bioanalyzer. Libraries were built using the PerkinElmer NextFlex Rapid Directional RNA-Seq Kit 2.0 (PerkinElmer catalog # NOVA-5198-02) according to the manufacturer’s instructions and sequenced with a NovaSeq 6000 (SE75).

### Assay for transposase-accessible chromatin by sequencing (ATAC-seq)

ATAC-seq sample preparation was performed as described previously^48^. In brief, organoids/tumoroids were extracted from Matrigel using GCDR, pretreated with 200 U/ml DNase (Worthington) for 30 min at 37°C, then washed 3x in cold PBS. After counting, 50 thousand viable cells were resuspended in cold lysis buffer (10 mM Tris-HCl pH 7.4, 10 mM NaCl, 3 mM MgCl_2_, 0.1% NP40, 0.1% Tween-20, and 0.01% Digitonin) and incubated on ice for 3 minutes. After washing, nuclei were resuspended in 50 µl of transposition reaction mix (nuclease-free water (22.5 µl), TD buffer (25 µl), Tn5 Transposase (2.5 µl)) and incubated for 30 min at 37°C. The transposed DNA fragments were then purified using MinElute PCR Purification Kit (Qiagen), barcoded with Nextera dual indexes from Illumina and PCR amplified for up to 11 cycles. The amplified libraries were purified using Qiagen MinElute PCR Purification Kit (Qiagen cat# 28004) and fragment sizes were checked on a Bioanalyzer station using D1000 DNA High Sensitivity Chips (Agilent cat# <u>5067-5585</u>). Libraries underwent size selection before a second round of quality check and quantification (Qubit). Normalized libraries were then pooled and sequenced on a Novaseq 6000 (Illumina).

### Sample preparation for single cell RNA-seq and single cell Multiome

Organoids/tumoroids were washed in cold basal media, resuspended in Cell Recovery Solution (Corning #76332-050) and incubated on a shaker for 15 minutes at 4°C. The pellets were dissociated into single cells using TrypLE and washed twice in basal medium before being resuspended in PBS containing 0.04% bovine serum albumin (BSA). Single cell preparations were counted and processed following the manufacturer’s instructions (single cell 3’ v3.1 protocol, 10x Genomics). In brief, single cells were resuspended in a master mix and loaded together with gel beads and partitioning oil into the chip to generate the gel bead-in-emulsion (GEM). The poly-A RNA from each GEM was retrotranscribed to cDNA, which contains an Illumina R1 primer sequence, a Unique Molecular Identifier (UMI), and the 10x barcode. The pooled barcoded cDNA was then purified with Silane DynaBeads, amplified by PCR and the appropriately sized fragments were selected for subsequent library construction. During the library construction Illumina R2 primer sequence, paired-end constructs with P5 and P7 sequences, and a sample index were added.

For the multiome experiment, following nuclei dissociation, scATAC-seq targeting 10,000 cells was performed using Chromium Next GEM Single Cell ATAC Library & Gel Bead Kit v.1.1 (10x Genomics, cat#1000175) and snRNA-seq targeting 10,000 cells was performed using Chromium Next GEM Single Cell 3′ Reagent Kits v.3.1(10x Genomics, cat#1000121). Samples were sequenced on a NovaSeq 6000 (Illumina).

### Bulk RNA-seq sample processing and analysis (alignment and quality control)

Quality control has been performed using FastQC (v0.11.8, RRID:SCR_014583) (Andrews 2010). Trim Galore! (v0.6.6, RRID:SCR_011847) was used to trim the adapter sequences with a quality threshold of 20 (Krueger 2012). The mouse genome reference, GRCm38.p6 and GENCODE release M25, was used as the transcriptome reference (RRID:SCR_014966). The alignment was performed using STAR aligner (v2.7.5b, RRID:SCR_004463) (Dobin *et al.* 2013). Gene level read counts were obtained using Salmon (v1.2.1, RRID:SCR_017036) for all libraries (Patro *et al.* 2017). Sample normalization was carried out using the median-ratios normalization method from DESeq2 R (v1.30.1, RRID:SCR_015687), and differential expression analysis was performed using DESeq2 (Love *et al.* 2014). Genes with less than 5 reads in total across all samples were filtered as inactive genes. A gene was considered differentially expressed if the Benjamini-Hochberg adjusted p-value is less than 0.01 and the absolute log2 fold change is greater than 1. To quantify and overlap differentially expressed genes from multiple comparisons, bar plots and Venn diagrams were plotted using ggplot2 (v3.3.5, RRID:SCR_014601) and VennDiagram (v1.6.20, RRID:SCR_002414), respectively (Wickham H, 2016; Chen H & Boutros P.C., 2011). Gene set enrichment analysis was performed on gene lists sorted by -log10 of p-value * log2 fold change using clusterProfiler (v4.2.2, RRID:SCR_016884), and enrichment of selected terms was visualized using gseaplot2 from enrichplot (v1.14.2) and bubble plots were generated by ggplot2 (Wu *et al*. 2021; Yu G 2023). R (v.4.1.0, RRID:SCR_001905) was used to perform all bioinformatics analyses.

### Bulk ATAC-Seq sample processing and analysis (preprocessing and peak calling)

Quality control has been performed using FastQC (v0.11.8, RRID:SCR_014583). Trim Galore! (v0.6.6, RRID:SCR_011847) was used to trim the adapter sequences with a quality threshold of 20. For each individual sample, paired-end 75-bp reads were aligned to the mouse reference genome (GRCm38.p6 and GENCODE release M25, RRID:SCR_014966) using Bowtie2 (v2.1.0, RRID:SCR_016368) with parameters –q –X 2000. Reads were sorted using SAMtools (v1.11, RRID:SCR_002105), and mitochondrial alignments were removed, keeping only reads with MAPQ 30 for downstream analysis (Langmead B *et al*. 2013; Danecek *et al*. 2021). Picard (v2.2.4, RRID:SCR_006525) was used to remove duplicates (Picard Toolkit 2019). Post-filtering BAM files for samples in the same dataset were merged using SAMtool merge function, followed by peak calling using macs (v2.1.0, RRID:SCR_013291) with parameters --nomodel --nolambda – keepdup all –slocal 10000 and -q 0.001 (Zhang *et al*. 2008). Quantification of reads in significant peaks for all samples was performed using BedTools multicov (v2.29.2, RRID:SCR_006646) (Quinlan and Hall, 2010). Differential peak analysis was performed using DESeq2 (adjusted p-value < 0.01 and absolute log2 fold change >= 1.5, RRID:SCR_015687). Coverage tracks (Bigwig files) were generated from BAM files for individual replicates and for each condition (BAM file of replicates from the same condition were merged) using deepTools (v3.2.1, RRID:SCR_016366) bamCoverage with parameters --normalizeUsingRPKM --binsize10, and replicates from the same condition (Ramírez F et al., 2016). Hierarchical clustering of the average accessibility in significantly differential accessible regions of each condition was performed using the ‘hclust’ function from R and visualized using the deepTools plotHeatmap function.

### Motif Enrichment and transcription factor footprinting analysis

Motif analysis was performed using findMotifsGenome.pl from HOMER (v4.10, RRID:SCR_010881) on the summit of peaks with parameter -size 200 (Heinz S *et al*. 2010). Footprinting analysis was performed using Tobias (v0.13.2) on aggregated BAM files per condition (Bentsen, M. *et al*. 2020). ATACorrect function was applied to correct Tn5 insertion bias, and then ScoreBigwig was implemented to calculate transcription factor (TF) binding scores in peak regions. Differential binding fold change and p-value for each TF was calculated using the BINDetect function, and results of selected TF were visualized using PlotAggregate function. Elbow plot highlighting the top differential binding TFs were generated using the ggplot2, which shows the -log10 of pvalue * the log2 fold change of binding score.

### TCGA bulk RNA-seq data processing

STAR raw gene counts of RNA-seq data were downloaded for TCGA-COAD (57 normal and 453 primary tumor samples) and TCGA-READ projects (13 normal and 163 primary tumor samples). Data from each project was normalized by median-ratios normalization, and the differential expression of genes between primary tumor and solid tissue normal was performed using DESeq2 (RRID:SCR_015687). Gene set enrichment analysis was performed on gene lists sorted by -log10 of p-value * log2 fold change using clusterProfiler (v4.2.2, RRID:SCR_016884), and enrichment of selected terms was visualized using gseaplot2 from enrichplot (v1.14.2). To correlate TCGA-COAD samples with mouse WT-RXRi and WT-DMSO samples, log CPM counts were generated for each dataset using the cpm function from edgeR (v3.36.0, RRID:SCR_012802) package (Robinson MD *et al*. 2010; McCarthy DJ *et al*. 2012; Chen Y *et al*. 2016). Average expression of each gene in normal and primary tumor conditions was calculated for TCGA project. Log CPM of TCGA samples and of the mouse samples were merged, and top 2000 most varied genes in terms of the log CPM values were selected, and Spearman correlations were calculated between samples on the selected genes.

### Public human bulk RNA-seq data processing

Human intestinal RNA-Seq FPKM data was retrieved from https://github.com/hilldr/Finkbeiner_StemCellReports2015/tree/master/DATA. To assess the similarity of expression profiles between GI track sites and the normal and tumor TCGA-COAD samples, FPKM version of the TCGA-COAD data was retrieved from TCGA data portal and was merged with the human intestinal FPKM data. Batch correction was applied using removeBatchEffect from limma R package (v3.50.1, RRID:SCR_010943), and prcomp was used to perform principal components analysis (Ritchie ME *et al*. 2015, RRID:SCR_014676). Differential expression analysis was performed using eBayes with trend = FALSE for the GI track data, and the top 300 up-regulated and down-regulated genes between adult intestine and fetal intestine were identified and used as signature for GSEA.

### Single cell RNA-seq data processing of mouse organoids

Sequenced FASTQ files were aligned, filtered, barcoded, and UMI counted using Cell Ranger Chromium Single Cell RNA-seq version 6.0.1, by 10X Genomics with Cell Ranger, mm10 (version 2020-A) as the mouse genome reference (RRID:SCR_017344). Each dataset was filtered to retain cells with ≥ 1000 UMIs, ≥400 genes expressed, and <10% of the reads mapping to the mitochondrial genome. UMI counts were then normalized so that each cell had a total of 10,000 UMIs across all genes and these normalized counts were log-transformed with a pseudocount of 1 using the “LogNormalize” function in the Seurat package. The top 2000 most highly variable genes were identified using the “vst” selection method of “FindVariableFeatures” function and counts were scaled using the “ScaleData” function. Datasets were processed using the Seurat package (version 4.0.3, RRID:SCR_016341) (Hao Y *et al*. 2021).

Principal component analysis was performed using the top 2000 highly variable features (“RunPCA” function) and the top 30 principal components were used in the downstream analysis. Datasets were integrated by using the “RunHarmony” function in the harmony package (version 0.1.0) (Korsunsky *et al*. 2019). K-Nearest Neighbor graphs were obtained by using the “FindNeighbors” function and the UMAPs were obtained by the “RunUMAP” function. The Louvain algorithm was used to cluster cells based on expression similarity. The resolution was set at 1.6 for optimal clustering.

Differential markers for each cluster were identified using the Wilcox test (“FindAllMarkers” function) with adjusted p-value < 0.01, absolute log2 fold change > 0.25, and minimum 10% of cells expressing the gene in both comparison groups using 1000 random cells to represent each cluster. Gene expression signature scores were calculated using the AddModuleScore function from Seurat package. Cells were annotated using the gene signatures for the cell types of interest by assigning each cell to the cell type with maximum module score. For each cell type, the contribution of each genotype is visualized as the percentage of cells in each cell type within each genotype.

RNA velocity analysis was performed using the scvelo Python package (version 0.2.5, RRID:SCR_018168) (Bergen *et al*. 2020). The unspliced and spliced count matrices were generated from Cell Ranger outputs using the run10x function from velocyto package (version 0.17.17, RRID:SCR_018167) (La Manno *et al*. 2018). For each sample and Harmony integrated dataset, Seurat object was converted into anndata (version 0.8.0, RRID:SCR_018209) (Isaac *et al*. 2021) and merged with the unspliced and spliced count matrices using scanpy (v1.9.3, RRID:SCR_018139) (Wolf *et al*. 2018). Moments for velocity estimation were calculated using 30 principal components and 30 neighbors. RNA velocity was estimated using a stochastic model with default parameters using scvelo. Cell type labels from the Harmony integrated dataset were projected to individual datasets and velocity streams were visualized.

### Single cell RNA-seq data processing of published human datasets

Raw count matrices for Lee *et al*. (Lee et al. 2020) and Pelka *et al*. (Pelka et al. 2021) were obtained from GEO with accession numbers GSE132465 and GSE178341, respectively. Seurat objects were created using raw count matrices. The Lee *et al*. dataset was filtered to keep cells from the patients with matching normal and tumor tissue. Both datasets were subset to only include the epithelial cells based on the available cell type annotations. UMI counts were then normalized so that each cell had a total of 10,000 UMIs across all genes and these normalized counts were log-transformed with a pseudocount of 1 using the “LogNormalize” function in the Seurat package. The top 2000 most highly variable genes were identified using the “vst” selection method of “FindVariableFeatures” function and counts were scaled using the “ScaleData” function. Datasets were processed using the Seurat package (version 4.0.3, RRID:SCR_016341) (Hao Y *et al*. 2021). Principal component analysis was performed using the top 2000 highly variable features (“RunPCA” function) and the top 30 principal components were used in the downstream analysis. UMAPs were obtained by the “RunUMAP” function. Tissue type-specific gene signatures from Lukonin *et al*. (Lukonin *et al*. 2020) were used for assigning each cell to a tissue type. Gene expression signature scores were calculated using AddModuleScore function from the Seurat package. Cells were annotated using the gene signatures by assigning each cell to the tissue type with maximum module score. For each sample, the contribution of each tissue type was visualized as the percentage of cells in each tissue type within each sample.

To identify human oncofetal gene signature, we created a Seurat object using the TPM counts from Gao et al (Gao *et al*., 2018). This Seurat object is filtered to keep fetal epithelial and adult cells from 6-and 7-week-old embryos. Pseudobulk samples are created by aggregating counts using Muscat (v1.12.1, Crowell *et al*., 2020). DESeq2 is performed to compare fetal and adult samples to obtain a fetal gene signature (adjusted p-value < 0.01 and absolute log_2_ fold change > 1). This human fetal gene list is further filtered to only keep the human orthologs of neoplastic cluster (c22) differentially expressed genes (adjusted p-value < 0.01 and absolute log_2_ fold change > 1) to obtain the human oncofetal gene signature.

### Single nucleus RNA-seq data processing

Sequenced FASTQ files were aligned, filtered, barcoded, and UMI counted using Cell Ranger ARC version 2.0.0, by 10X Genomics. mm10 database (version 2020-A) was used as the mouse genome reference. Each dataset was filtered to retain cells with ≥ 1000 UMIs, ≥400 genes expressed, and <10% of the reads mapping to the mitochondrial genome. UMI counts were then normalized so that each cell had a total of 10,000 UMIs across all genes and these normalized counts were log-transformed with a pseudocount of 1 using the “LogNormalize” function in the Seurat package. The top 2000 most highly variable genes were identified using the “vst” selection method of “FindVariableFeatures” function and counts were scaled using the “ScaleData” function. Datasets were processed using the Seurat package (version 4.0.3, RRID:SCR_016341) (Hao Y *et al*. 2021).

Principal component analysis was performed using the top 2000 highly variable features (“RunPCA” function) and the top 30 principal components were used in the downstream analysis. K-Nearest Neighbor graphs were obtained by using the “FindNeighbors” function whereas the UMAPs were obtained by the “RunUMAP” function. The Louvain algorithm was used to cluster cells based on expression similarity. The resolution was set at 1.6 for optimal clustering.

### Single-cell ATAC-seq data processing and motif analysis

Sequenced FASTQ files were aligned, filtered, barcoded, and UMI counted using Cell Ranger ARC version 2.0.0, by 10X Genomics. mm10 database (version 2020-A) was used as the mouse genome reference. Peaks were called using ‘CallPeaks’, the MACS2 wrapper function, in Signac (v1.3.0, RRID:SCR_021158) (Stuart, T *et al*. 2021). Cells were filtered to retain only those with total number of fragments in peaks > 1000 and < 50000, percentage of reads aligned in peak regions > 20%, blacklist ratio < 0.002, nucleosome signal < 1.5 and the TSS enrichment score > 2. The blacklist ratio was calculated as the number of reads aligned to blacklist region divided by the total number of reads. The nucleosome signal was calculated using the ‘NucleosomeSignal’ function in Signac, and the TSS enrichment score was calculated by ‘TSSEnrichment’. Latent semantic indexing (LSI) was then applied to the top 95% of peaks in terms of their variability, which combines the term frequency-inverse document frequency (TFIDF) used for normalization and singular-value decomposition (SVD) used for dimensional reduction. The first LSI component was excluded from all downstream analysis as it highly correlated with sequencing depth. De-novo clustering at resolution 1.6 was performed by using ‘FindNeighbors’ and FindClusters functions in Signac. Motif position frequency matrices were retrieved from JASPAR 2020 database (RRID:SCR_003030) and were added to the snATAC data in Signac (Baranasic D *et al*. 2022). ChromVAR motif analysis was performed using the RunChromVAR function in Signac, and results of selected transcription factor were visualized on UMAP using FeaturePlot function (Schep, A. N *et al*. 2017).

### Integration of multiome data and defining meta-clusters

Signature module scores were calculated on snRNA expression data using the AddModuleScore function in Seurat (v 4.0.3, RRID:SCR_016341), and the module scores were projected to, and visualized on, a snATAC UMAP (Hao Y *et al*. 2021). Average fetal module scores and average stem module scores for each of the snATAC clusters were calculated by averaging the signature scores of all cells in each cluster and were used to define four meta-clusters (none, dual, oncofetal and stem). The median of average cluster signature scores were used as thresholds, and clusters with signature scores above the median were defined as positive for that signature type. Clusters that were positive for both signatures formed the “dual” meta-clusters, and clusters that were negative for both signatures formed the “none” meta-cluster.

### Statistics & reproducibility

All experiments were reproduced at least 2-3 times. For *in vivo* experiments, data are represented as mean ± s.e.m. None of the experiments were blinded and no statistical methods were used to pre-determine sample size. The number of animals/injections was estimated based on the variability in tumor take rate and growth and is indicated in the respective figure legends. Animals were randomized among treatment groups when tumors reached a mean volume of approximately 360mm^3^.

R (v.4.1.0, RRID:SCR_001905) was used for all statistical analysis and visualization (R Core Team, 2023). All boxplots show the median and the inter quartile range (IQR) of the underlying data with whiskers extending to ± 1.5 IQR from boxes. All outliers are shown unless stated otherwise. All figures were generated using ggplot2 (v3.3.5, RRID:SCR_014601), ggpubr (v0.4.0, RRID:SCR_021139) and patchwork (v3.0.1) (Kassambara A., 2023, Wickham H., 2016, Pedersen T., 2022). We used the Wilcoxon test for statistical testing unless stated otherwise. Significance levels are defined as follows: n.s: p > 0.05, ∗p ≤ 0.05, ∗∗p ≤ 0.01, ∗∗∗p ≤ 0.001 and ∗∗∗∗p ≤ 0.0001.

---

**Extended data Fig. 1:**
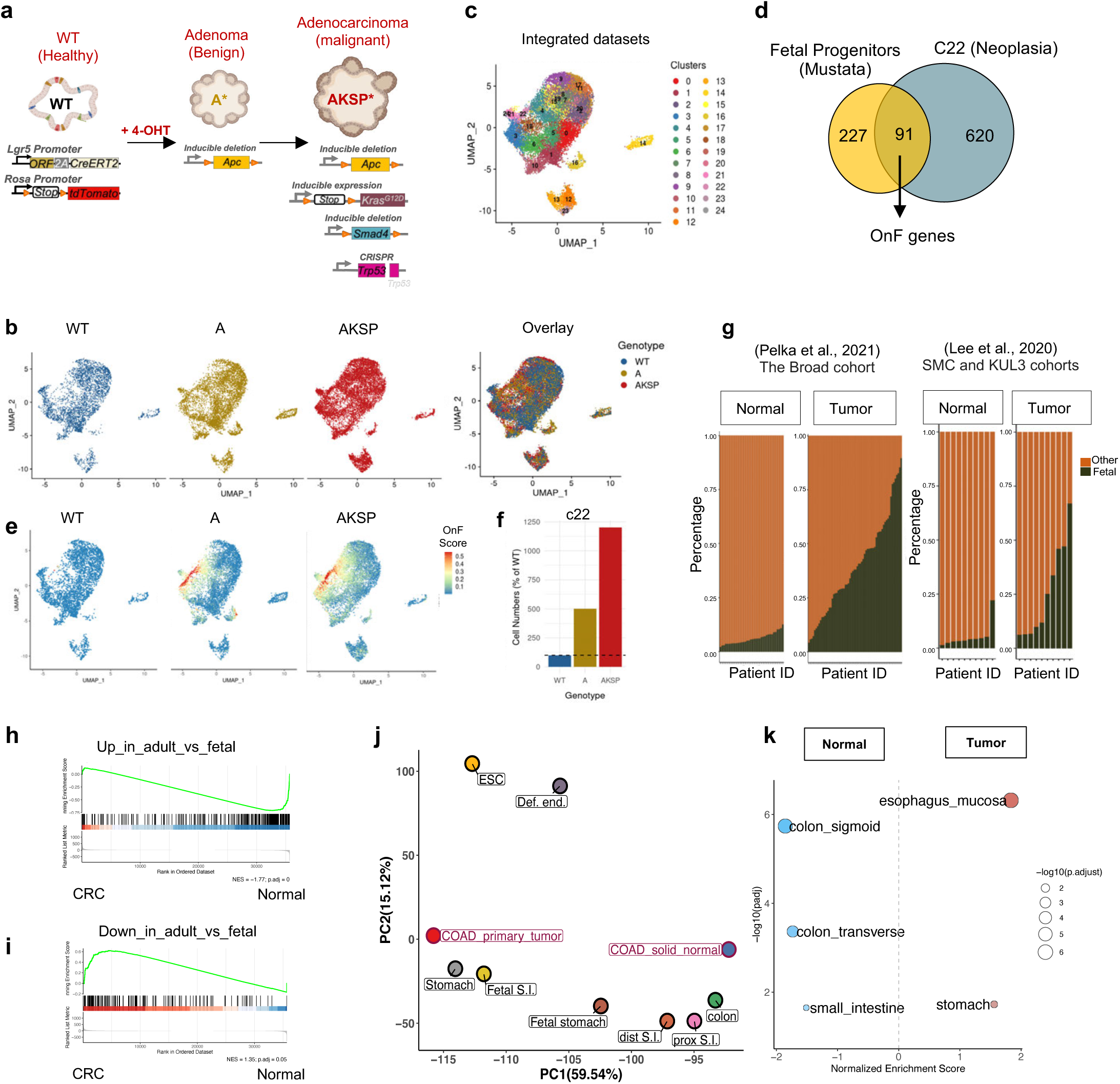
Oncofetal reprogramming triggers lineage plasticity in CRC. **a,** Schematics of the genetic approach used to generate isogenic mouse organoid models that mimic the adenoma to adenocarcinoma sequence. **b,** Cells from indicated organoid models are color coded and visualized separately (left) and overlayed (right) in the integrated UMAP embedding as in Fig 1b**. c,** Louvain clustering of cells based on gene expression from organoids models indicated in **(a, b).** Unsupervised clusters are overlaid on the UMAP and indicated by different colors. **d,** Venn diagram highlighting the overlap between known intestinal fetal markers (add reference), and the tumor-specific cluster 22 (c22). The shared genes are used to define the oncofetal (OnF) signature. **e,** Enrichment of the OnF signature across the CRC malignancy continuum depicted by the indicated organoid models. **f,** Proportion of cells contributing to c22 in WT, A and AKSP organoids shown as percent enrichment over proportion in WT organoids. **g,** Proportion of normal and tumor cells with positive enrichment of OnF signature in the Broad (left) and SMC and KUL3 (right) cohorts. **h, i,** Gene Set Enrichment Analysis (GSEA) of adult **(h)** and fetal **(i)** human intestine markers in the COAD/TCGA dataset (n=41 and 455 for normal and CRC samples, respectively). **j,** Principal Component Analysis (PCA) of integrated bulk RNA-seq data from the indicated developmental stages of the human gut tube, definitive endoderm (Def. end), embryonic stem cells (ESC), normal, and CRC samples from the TCGA/COAD dataset. **k,** Enrichment analysis of the indicated tissue-specific signatures in tumor vs. normal specimen^ from the COAD/TCGA dataset. Red and blue indicate positive and negative normalized enrichment scores (NES), respectively. Bubble size indicates the negative of log _w_ adjusted p-value.

**Extended data Fig. 2:**
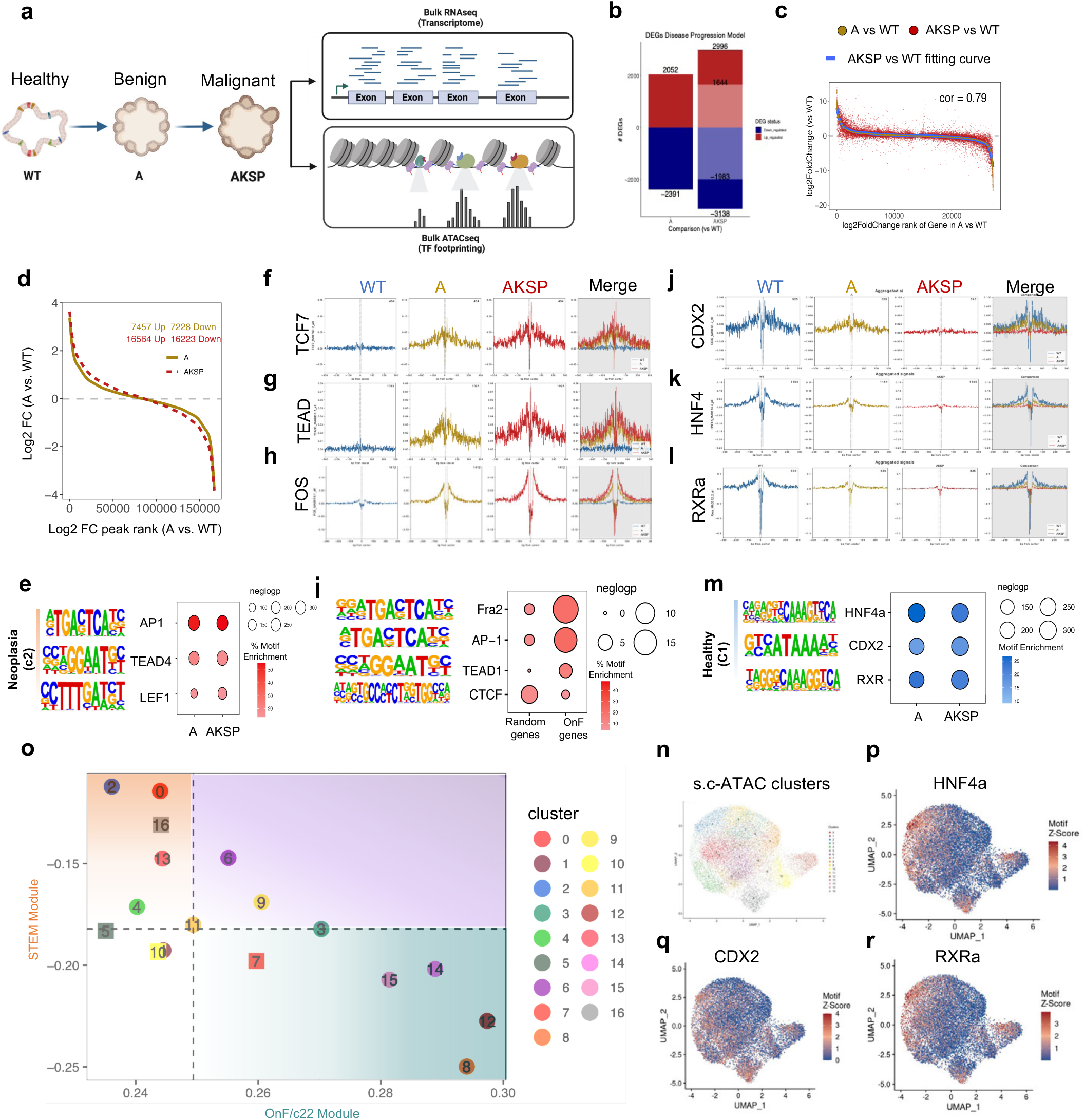
A multi-omics approach elucidates the determinants of cell state dynamics in CRC. **a,** Schematics of the experimental strategy and genetic models used. **b,** Differentially expressed genes (DEGs) in A vs. WT and AKSP vs. WT. Shared DEGs are indicated with faded colors. **c,** Correlation plot of DEGs in A vs. WT (yellow dots) and AKSP vs. WT (red dots and fitted blue curve) in the upper panel. **d,** Comparison of differentially accessible regions (DARs;log2 fold change) in AKSP vs. WT (dashed line) and A vs. WT as a reference (solid line). geom_smooth function from ggplot2 package is used to fit the curve to fold changes in both comparisons. **e,** HOMER motif analysis for the indicated TFs, using differentially accessible regions (DARs) from cluster c2 in Fig. 2c. **f, g, h,** TF occupancy strength presented as aggregated footprinting plot matrix for the indicated TFs in WT, A and AKSP organoids. Plots are centered around binding motifs**. i,** HOMER motif analysis for the indicated TFs, using DARs in promoter regions (TSS +/-1kb) of OnF genes or random regions. **j, k, l,** Same analysis as in **f, g, h** for a different set of TFs. **m,** HOMER motif analysis for the indicated TFs, using DARs from cluster c1 in Fig. 2c. **n,** UMAP embedding of scATAC-seq cells from AKSP tumoroids, colored by unsupervised clusters at resolution 1.6. **o,** Scatter plot showing the distribution of scATAC-seq cell clusters based on their OnF (x-axis) and stem cell (y-axis) enrichment scores. Four distinct meta-clusters have been defined accordingly. **p, q, r,** TF motif activity in scATAC-seq cells determined by chromVAR deviation scores for the indicated TFs.

**Extended data Fig. 3:**
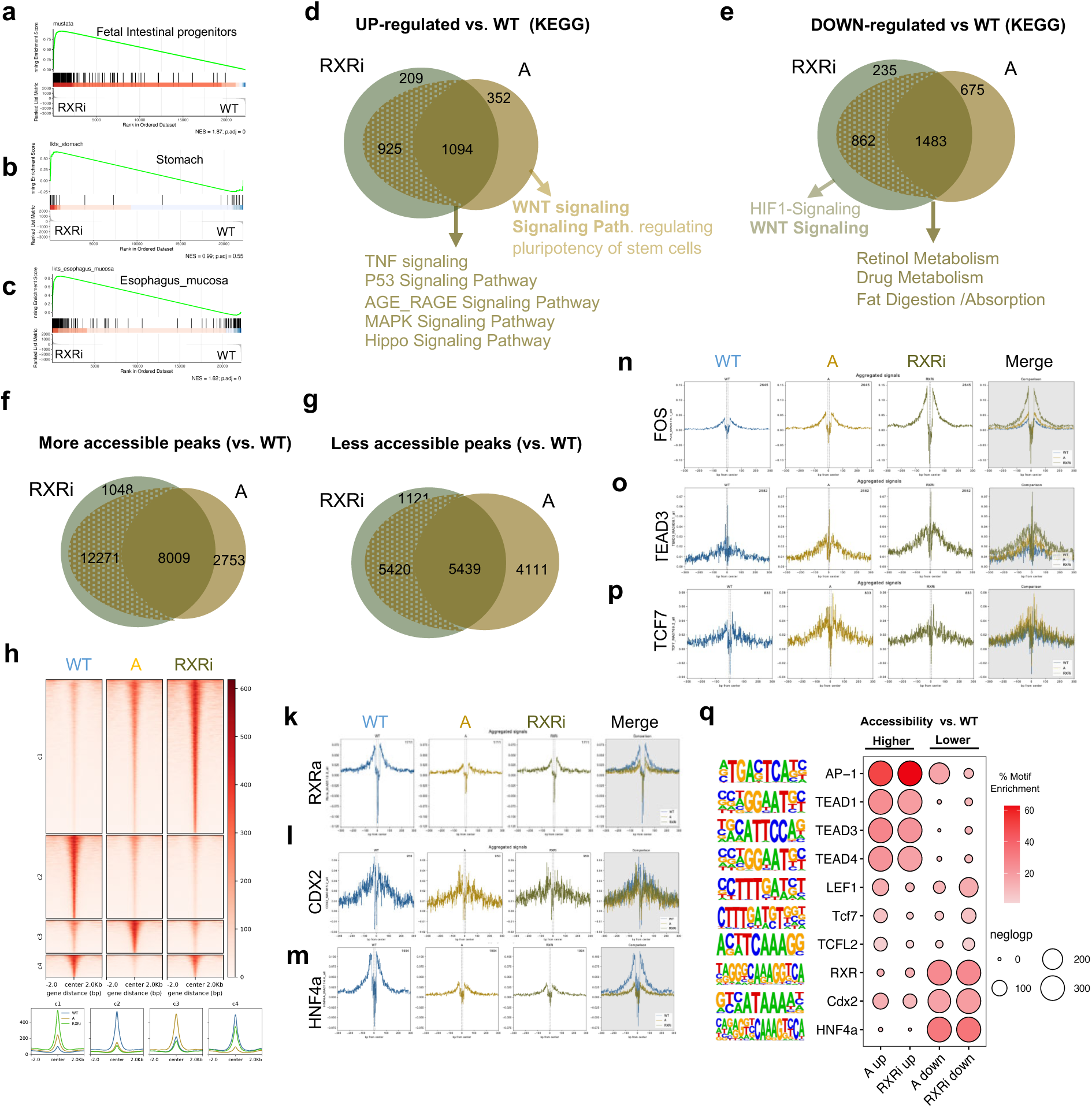
RXR inhibition phenocopies APC depletion excluding WNT activation. **a, b, c,** GSEA of the indicated transcriptional signatures in RXRi vs. WT organoids. **d, e,** Venn diagrams depicting the overlap of up-regulated **(d)** and down-regulated **(e)** genes following RXR inhibition and APC depletion in WT organoids. Punctuated areas represent changes in the same direction that did not make the cutoff of 2-fold change. KEGG pathway analysis highlights a common regulation of most molecular pathways in both models, except for WNT. **f, g,** Venn diagrams depicting the overlap of differentially accessible regions in the RXRi and A organoid models. **h,** Heatmap of ATAC-seq signal in WT, A and RXRi organoids at differentially accessible regions (upper panel). Comparison of average ATAC-seq signal profile in the indicated organoid lines within a +/-2Kb region around the peak center in the different clusters (lower panel). **k, l, m, n, o, p,** TF occupancy strength presented as aggregated footprinting plot matrix for the indicated TFs in WT, A and RXRi organoids. Plots are centered around binding motifs. **q**, HOMER motif analysis for the indicated TFs, using differentially accessible regions in A and RXRi models compared to WT organoids.

**Extended data Fig. 4:**
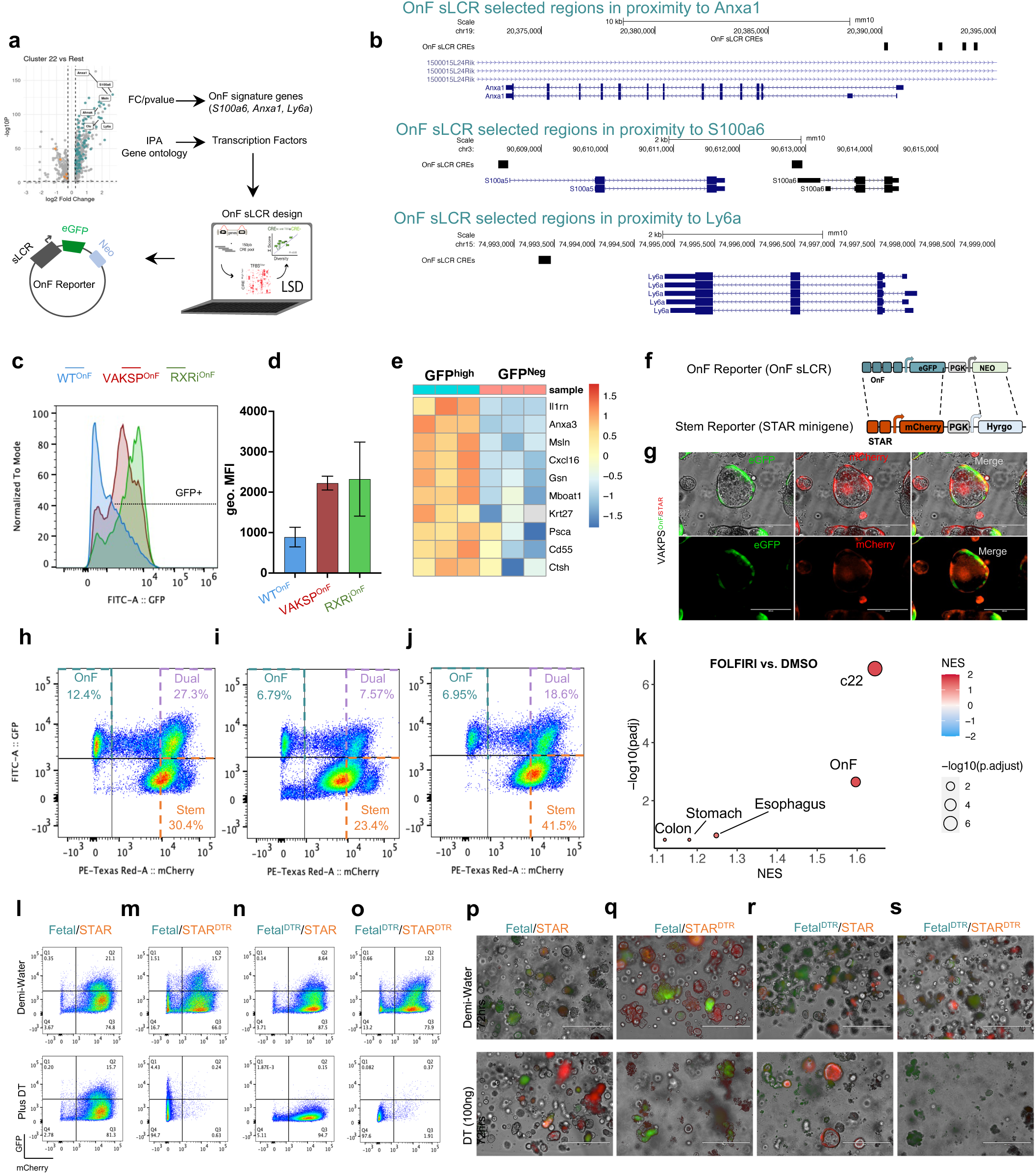
Design and functional validation of the oncofetal phenotypic reporter. **a,** Schematic representation of the oncofetal (OnF) synthetic locus control regions (sLCR) generation from gene expression data **b,** UCSC Genome Browser view of the regions used as cis­regulators elements in the OnF sLCR. **c,** Normalized percent distribution of OnF-GFP+ cells and eGFP fluorescence intensity in the indicated organoid/tumoroid models determined by FACS. **d,** Quantification of the eGFP geometric Mean Fluorescence Insensitivity (MFI) in organoid models from panel **(c)**. **e,** Heatmap of OnF gene expression in GFP^high^ and GFP^Neg^ cells sorted form VAKSP^OnF^ tumoroids. **f,** Schematic representation of substitution of the OnF sLCR with the STAR minigene. eGFP and the neomycin resistance cassette were replaced with mCherry and a hygromycin resistance cassette, respectively. **g,** Representative fluorescence microscopy images of VAKSP tumoroids co-expressing the OnF and stem-cell specific (STAR) reporters (VAKSP^OnF/STAR^). **h, i, j,** Representative pseudocolor plots of flow cytometry analysis of VAKSP^OnF/STAR^ tumoroids co­expressing the oncofetal (OnF-GFP, Y axis) and stem (STAR-mCherry, X axis) phenotypic reporters at different times of culture. **k,** Enrichment analysis of the indicated gene-set signatures in VAKSP tumoroids treated with FOLFIRI for 3 days compared to DMSO control. Red and blue indicate positive and negative normalized enrichment scores (NES), respectively. Bubble size indicates the negative log_10_ p-value15**, m, n, o,** Representative pseudocolor plots of flow cytometry analysis of the indicated tumoroid lines following treatment with demi-water or DT for 3 days. **p, q, r, s,** Representative fluorescence microscopy images of the indicated VAKSP tumoroid models treated with demi-water or DT for 3 days.

